# Laminar signal extraction over extended cortical areas by means of a spatial GLM

**DOI:** 10.1101/285544

**Authors:** Tim van Mourik, Jan PJM van der Eerden, Pierre-Louis Bazin, David G Norris

**Affiliations:** Radboud University Nijmegen, Donders Institute for Brain, Cognition and Behaviour, Nijmegen, The Netherlands; Spinoza Centre for Neuroimaging, Amsterdam, the Netherlands; Netherlands Institute for Neuroscience, Amsterdam; Max Planck institute for Human Cognitive and Brain Sciences, Leipzig, Germany; Erwin L. Hahn Institute for Magnetic Resonance Imaging, Essen, Germany

## Abstract

There is converging evidence that distinct neuronal processes leave distinguishable footprints in the laminar BOLD response. However, even though the achievable spatial resolution in functional MRI has much improved over the years, it is still challenging to separate signals arising from different cortical layers. In this work, we propose a new method to extract laminar signals. We use a spatial General Linear Model in combination with the equivolume principle of cortical layers to unmix laminar signals instead of interpolating through and integrating over a cortical area: thus reducing partial volume effects. Not only do we provide a mathematical framework for extracting laminar signals with a spatial GLM, we also illustrate that the best case scenarios of existing methods can be seen as special cases within the same framework. By means of simulation, we show that this approach has a sharper point spread function, providing better signal localisation. We further assess the partial volume contamination in cortical profiles from high resolution human ex vivo and in vivo structural data, and provide a full account of the benefits and potential caveats. We eschew here any attempt to validate the spatial GLM on the basis of fMRI data as a generally accepted ground-truth pattern of laminar activation does not currently exist. This approach is flexible in terms of the number of layers and their respective thickness, and naturally integrates spatial regularisation along the cortex, while preserving laminar specificity. Care must be taken, however, as this procedure of unmixing is susceptible to sources of noise in the data or inaccuracies in the laminar segmentation.

## 1 Introduction

With functional Magnetic Resonance Imaging (fMRI) neuronal activity in the brain is measured indirectly via the Blood Oxygen Level Dependent (BOLD) response. With the emergence of higher static magnetic fields, more powerful acquisition sequences and better analysis tools, the location of the activation can be pinpointed more precisely. The attainable spatial resolution can be so high that voxels are smaller than the thickness of the cerebral cortex. These improvements have made it possible to investigate specific cortical layers with fMRI. Typically the human cerebral cortex consists of sixcytoarchitectonic layers [1]. Layer IV is commonly associated with receiving feedforward input from Layer III from lower cortical areas or from the thalamus [2], while Layers II-III and VI are implicated in receiving downward information flow (feedback) [3], which often originates from layer V. Layer I is thin and sparsely populated with neurons and will probably remain elusive to laminar fMRI.

It is clear that there may be a lot of information about laminar processing in fMRI measures. The BOLD signal has convincingly been shown to have a laminar origins in the rat motor- and somatosensory cortices [4]. Further tight spatial coupling has been demonstrated of blood flow and dilation of arterioles of layer II/III and orientation tuning in the cat visual cortex [5]. And in line with previous depth-dependent electrode recordings, the BOLD response that uniquely reflects trial by trial variance in the alpha and gamma bands was recently shown to be consistent with infra- and supra-granular origins of these oscillations [6]. While the details of the neurovascular coupling are still unknown [7], the cortical BOLD response has been modelled as a function of depth and could potentially even be deconvolved to get a better estimate of the origin of cortical activation [8]. The work of Scheeringa et al. suggests that the laminar BOLD response as measured in humans [9–12, e.g.] contains distinguishable laminar responses. Recent advances in vascular space occupancy (VASO) techniques have shown cerebral blood volume (CBV) measurements to be laminarly specific as well [13]. If this is indeed the case, laminar fMRI could give us the means of measuring directional communication between brain regions. For this reason, extracting reliable and meaningful layer specific time courses in humans has been recognised as essential to get a better understanding of the nature of computations that are performed by the brain [14, 15].

Also from a neuroanatomical perspective, the cyto- and myeloarchitectonic layers play a central role, but they have classically been examined through invasive histology. Developments in MRI have allowed more detailed non-invasive investigation of the cortical anatomy [16]. This has further been related to functional studies [9, 17], allowed for more detailed anatomical parcellations [18, 19], and may even be linked to developmental changes [20]. Ultimately, detailed laminar anatomical investigations may yield biomarkers for pathological markers [16].

Hitherto little attention has been paid to the question of how to extract laminar signals from high spatial resolution fMRI data. Voxels are sometimes manually classified to be part of layers at different cortical depths [11, 21, 22, e.g.]. Other attempts included drawing lines perpendicular to the surface and interpolating the volume, either manually [9], or using a cortical mesh reconstruction [10, 23, 24, e.g.]. The variation in the distribution of the histological layers over cortical depth in gyri and sulci was identified as a challenge for laminar fMRI [25]. This is why several studies chose to analyse straight pieces of cortex only [9, 22, 24]. The way that the layer thickness varies over the cortex relates to the curvature and was found to behave according to an equivolume principle [26, 27], which can be modelled by means of a level set framework [28], or equivalently with a surface based sampling algorithm [29].

Even if the cytoarchitectonic layer topography was known throughout the cortex, it would still be challenging to extract laminar signals. As the fMRI data will generally consist of cubic voxels, these voxels will almost certainly contain signal from several layers. Any kind of interpolation will lead to contamination from neighbouring layers. This effect is reduced with higher resolution, but the contamination effect in relation to the spatial resolution has never been quantified. The term ‘laminar resolution’ [30, 31] has been used to roughly mean sub-millimetre resolution. While it is certainly improbable to get laminar specific results at lower resolutions, the one millimetre threshold is arbitrary. Given that the cortex is on average 3 millimetres thick [32, 33], the resolution requirements may well change dependent on the cortical area considered and the layers of interest.

Here we propose a method to reliably extract time courses from a cortical area by using the framework of the General Linear Model (GLM). This offers a potential solution to the partial volume problem, for the situation in which a common laminar signal can be assumed over a number of voxels that is large compared to the number of layers. Instead of interpolating and integrating, we propose to decompose the layer signals by means of a spatial GLM. While in the limit of infinitesimal voxel volume all methods should yield the same result, our method aims to retrieve more accurate results at coarser resolutions. An added benefit is that the mathematical assumptions underlying the GLM are known and their validity may be tested within a data set. This work has previously been presented in abstract form [34]. A similar suggestion for a laminar mixture model was presented in abstract form by Polimeni et al. [35].

Herein we describe the theory and implementation of the spatial GLM. We explain in detail the pipeline for laminar data processing and the extraction of the laminar profile. In order to test the power of the spatial GLM, we employed a simple simulation to generate a curved model cortex which satisfies the equivolume principle. This allowed us to set a gold standard on which we could test our method and compare it with other laminar signal extraction methods. In addition, we validated our method using high resolution structural data in order to show that we could obtain a profile that preserves underlying anatomical structures. Lastly, we tested whether we could extract robust profiles across grey matter from structural scans. We anticipate that the main use of the spatial GLM will be in the extraction of functional time courses, and have already utilised this technique to detect layer specific feedback signals in human primary visual cortex [12, 36]. In their respective supplementary materials, comparisons can be found with existing methods. In the current work our emphasis is on giving a full description of this technique, and validating it in situations where a known ground truth can be postulated. We eschew here any attempt to validate the spatial GLM on the basis of fMRI data as a generally accepted ground-truth pattern of laminar activation does not currently exist.

## 2 Theory

## 2.1 GLM

The framework of the General Linear Model (GLM) is routinely used in fMRI for fitting voxel time courses to a temporal model [37]. The GLM framework can also be used spatially, as illustrated for example by a dual regression [38]. Here we propose to use a spatial GLM where an *n* × *k* design matrix **X** represents the layer volume distribution, i.e. the distribution of the *k* layers over the *n* voxels within a region of interest. Every row of **X** gives the distribution of a given voxel volume over the layers and every column (regressor) represents the volume of the corresponding layer across voxels. It is assumed that, within a region of interest, the layer signal is uniform. The regression of the voxel signals against the design matrix yields the layer signal. The crucial difference with the current cortical layer and profile modelling methods is that the GLM decomposes the voxel signals into the respective layer signals. In contrast, interpolation does not make an attempt at unmixing the signal and will be subject to partial volume contamination that will result in signal leakage between neighbouring layers.

For any chosen voxel grid, the design matrix **X** should be derived from the location of the layers. These layers are not necessarily identical to architectonic layers, but instead reflective of a measure for cortical depth. In general the layer depths are not precisely known. In the present work we estimate the layer distribution, and hence the spatial design matrix, using the level set method [27], explicitly described in section 2.2. Layer boundaries constructed with this method can be viewed as snapshots of a surface moving smoothly from the white matter boundary to the pial boundary. Since it is assumed that the underlying laminar signal is constant throughout a region of interest (ROI), the voxel signals of the ROI can be regressed against this design matrix, yielding the estimated laminar signals from the ROI.

A general linear model with a number of voxels *n* and number of time points *m* can be described as:

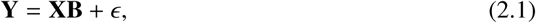

where, **Y**, size [*n* × *m*], represents a multivariate distribution that is being modelled by **X**, size [*n* × *k*], the laminar design matrix with *k* layers. The model is fit in order to obtain estimates 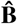, size [*k* ×*m*], and these estimates are chosen such, that the error term *E* is minimised. The columns in **X** (regressors) essentially represent the (fractional) presence of respective layer over all voxels. Note that a standard fMRI temporal regression would estimate **Y**^**T**^ instead of **Y**, such that **X** represents *temporal* regressors instead of *spatial* regressors.

For example an Ordinary Least Squares (OLS) estimation can be used for minimisation. This problem has a unique solution as long as **X** contains more rows (number of voxels) than columns (number of layers) and as long **X** is not rank deficient. **X** could be rank deficient when not every layer is represented in the design, or when the distribution of one layer is a linear combination of the others. The latter is highly unlikely but could occur when a high number of layers is computed and neighbouring layers occupy the same space, resulting in a collinear system. However, the matrix becomes increasingly ill-conditioned when the number of layers exceeds the number of voxels over the cortical thickness.

It should be noted that the mathematical framework of the GLM comes with (strong) assumptions. Each measurement is assumed to be independent and counts as a degree of freedom. The interpretation for MRI is that the intensity of a voxel should not be predictable based on observing its neighbours. This assumption is clearly violated in (f)MRI data, and as a result, the degrees of freedom (DoF) of the system will be overestimated. As a direct consequence, the standard error is underestimated giving erroneously small confidence intervals. We hence do not report or show single subject error estimates. Additionally, the mathematical framework assumes a linear mixture of an uniform effect (i.e. same layer intensity over space). This may pose severe problems in the presence of a bias field, intensity fluctuations across a layer, or structural variations such as cortical veins. Smart vein removal may hence be necessary for laminar fMRI [9, 39], before a spatial GLM is applied. Lastly, voxelwise errors should follow a specified distribution: Gaussian in case of General Linear Model, but in reality potentially following a different distribution (Rician, [40], or more complex when using parallel imaging techniques).

A variety of estimation methods can be used to solve this system of equations. While a simple way of obtaining layer intensities is an OLS estimation, it estimates 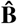 based on an *l*_2_-norm. Other regularisation techniques may be employed to improve the estimation. This can be done by either imposing constraints on the outcome, or by introducing prior knowledge. The first can be achieved by including entropy measures in the estimation such as a smoothness constraint (*λ*∥∇**B**∥_0_) or sparseness constraint (∥**B**∥_0_). However, these techniques bias the result in a certain direction. As this is undesirable for subsequent analyses, we do not further discuss them in this paper. A way of introducing prior knowledge into the estimation is by making assumptions about the covariance structure of the noise. OLS assumes that the voxelwise errors *ϵ* are independent, normally distributed with mean zero, and have constant variance: *ϵ*∼ *N*(0, *σ*^2^**I**). If a more general covariance matrix is assumed, *ϵ N*(0, Ω), estimation can be performed by Generalised Least Squares (GLS):

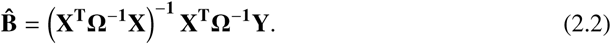

This requires an explicit description of the covariance matrix Ω. We propose to model this as a Gaussian as a function of the relative distance between voxels. The covariance is one when the distance is equal to zero, leading to a unity diagonal in Ω, and decreases rapidly for more distant voxels.

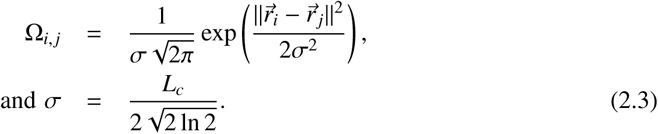

Here 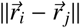 is the distance between voxel *i* and *j*. The standard deviation of the spatial Gaussian is *σ*. We here explore the impact of different correlation lengths, *L*_*c*_, defined as the FWHM for a Gaussian noise correlation. However, it would be possible to replace it with covariance matrices of different forms, e.g. the spatial point spread function of an EPI read-out.

### 2.2 Layer localisation

The most laborious part of the spatial regression is the construction of the layer volume distribution that acts as a design matrix. We used an in-house implementation of the level set method, originally proposed by [27].

Level set surfaces are a way to represent and manipulate surfaces in volumetric space. One way to obtain level sets is based on the signed distance function (SDF). This is the distance of a point to a given closed surface, for points enclosed by the surface the SDF is negative, for points outside the surface it is positive. Points with equal SDF define a surface in volume space. For such surfaces a mesh representation can be obtained that consists of vertices, edges and face, by means of a meshing algorithm (e.g. marching cubes [41]). The advantage of the SDF is that all computations can be performed in the same volume space as an MRI image. Lamination of the cortex can thus be represented in volume space. It is assumed that each laminar surface has a constant SDF. The level set is the set of corresponding SDF values, labelling regularly sampled layers between the white matter surface and the pial surface [27].

The way we calculated the cortical lamination differs slightly from Waehnert et al’s method [27]. Rather than computing the curvature at the surface, we compute it in eacg voxel based on the divergence of the Laplacian vector field. We consider vectors oriented along the direction of Laplacian streamlines from the white matter to the pial surface [42]. Based on the solution of the Laplace equation, we compute the gradient in the Fourier domain together with a Tukey window. This acts as a low-pass filter, such that the mesh representation is sufficiently smooth to calculate the mean curvature (half the surface divergence of the normal). As a result we can define a local curvature at each point of the cortex, which is then used to construct an equivolume level set of the different layers by means of the formula as given in Kleinnijenhuis et al. [29].

Having obtained the layer locations, the distribution of layers over voxels needs to be computed in order to create the layer volume distribution. This could be done by directly using a partial volume distribution as proposed by Koopmans et al. [23]: the average projection of a cube onto a line, for all possible orientations. Whereas Koopmans et al. [23] use it passively to estimate an effective resolution of a volume, it can be used actively to compute volumetric occupation of individual layers. This is directly represented by the integral of the partial volume distribution., where the integration limits are the distances provided by the level set. This can be made even more precise by taking into account the orientation of the voxel with respect to the cortex, instead of using the average of all possible orientations. Consider a voxel to be occupying a cubic space, that is intersected by a plane (the cortex) with normal 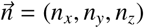 at distance *t*. The intersection area of the cube and the plane can then be calculated, for which an algorithm is described in Appendix B. From this, also the volumetric occupation of a layer over a voxel can readily be computed. This process is illustrated in Figure 1. This procedure easily generalises to multiple layers being present in a single voxel, as it corresponds to an intersection with multiple planes. The volumes for the respective layers are hence given by the integral of the intersection area from one plane to the next. The gradient estimate is voxel specific rather than layer specific (i.e. planes are assumed to be parallel) and the same intersection function is used. This is accurate as long as the voxel length is sufficiently small compared to the radius of curvature.

**Fig 1.**
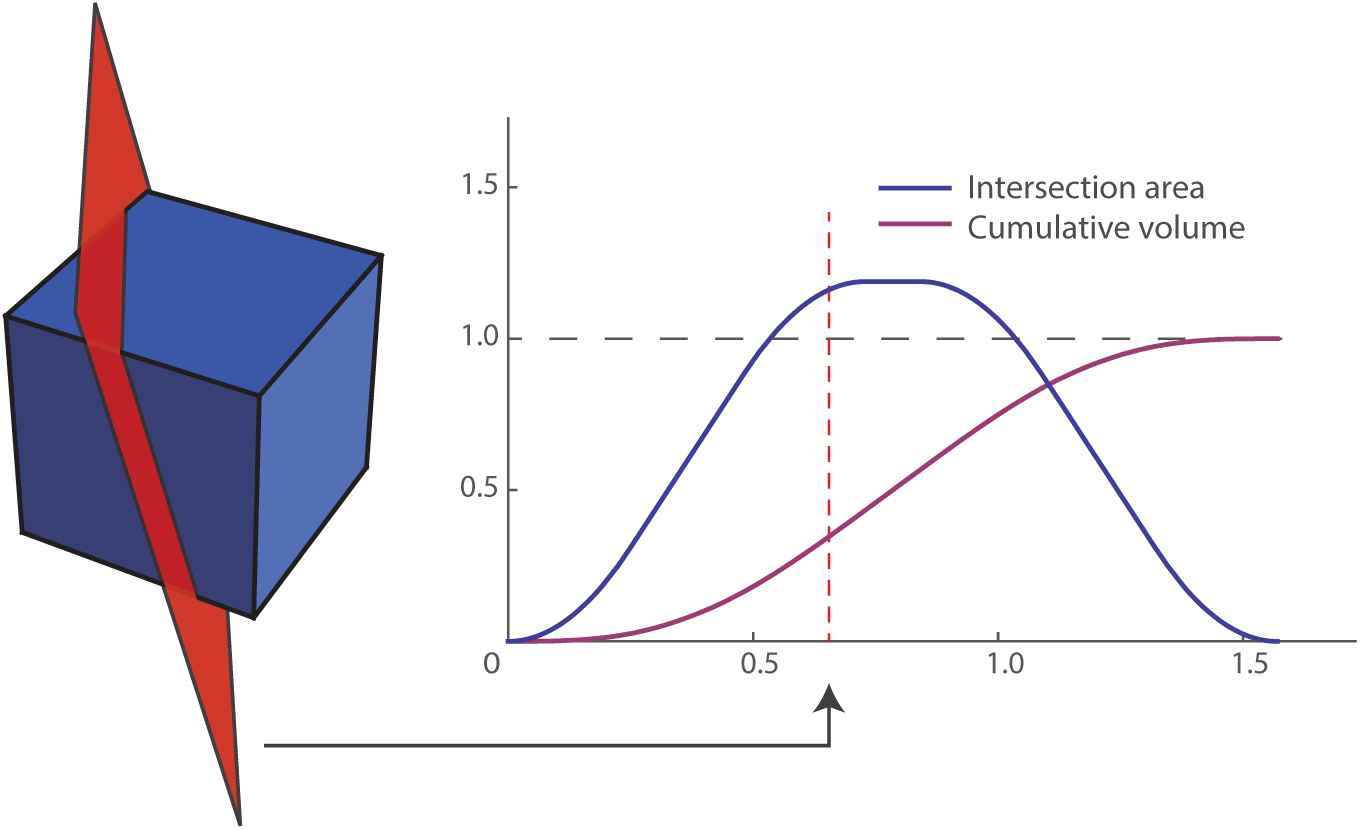
The plane of arbitrary normal 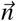 (here 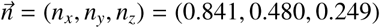) divides a unit voxel in two parts (red dashed line). As the plane moves in the direction of its the normal, the area of intersection varies, as indicated by the blue curve. The volume within the voxel on the left side of the plane is indicated by the cumulative volume and represented by the purple curve.

### 2.3 The laminar time course

Once the layer volume distribution is constructed, it can be applied to MRI data. For all given voxels within a region of interest, the voxel signal values represent our measurement data **Y**. The rows of the design matrix **X** give the fractions of the voxel volumes ascribed to the corresponding layer by the layer volume distribution. The layer estimates 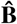 can be obtained by regression of **Y** against **X**, given covariance matrix Ω. In order to obtain a laminar time course from an ROI in fMRI time series, the regression can be performed sequentially for that ROI. Note that the unmixing matrix is independent of the temporal signal, so the regressor calculation needs only to be performed once.

### 2.4 Similarity to existing methods

Hitherto, two main methods of extracting laminar time courses have been used. In the first one, the cortical surface is represented by two triangular meshes, the white matter surface and the pial surface. A laminar profile is then obtained by drawing lines from points (vertices) on one surface to the other. The volume projected onto these lines gives a cortical profile. In computing this projection, the volume has to be sampled by means of some interpolation method. This approach has been used in a number of implementations [10, 23, 24]. The second method is a classification of each voxel to be in a given layer based on the single most likely layer per voxel. The signal is subsequently averaged over the region of interest [11, 21, 22]. Interestingly, all methods can be seen in the light of the same mathematical framework.

Interpolating a volume at different cortical depths across a part of the cortex effectively creates a weighting for all voxels with respect to the layers. The weighting in this procedure is based on a limited set of vertices that form the mesh. While it is not guaranteed that all voxels in the region of interest are equally represented, one could likely assume that in the limit of an infinite number of lines the result would be similar to our laminar design matrix **X**. The way in which the average is taken for all lines is then equivalent to a multiplication with the data, normalised with respect to the number of voxels:

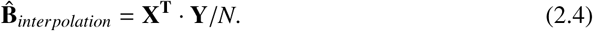

Here 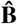, **X**, and **Y** are respectively the estimated layer signals, the weighting matrix, and the voxel signals and have the same dimensions as in Eq.2.1. *N* is the number of voxels. We argue that such multiplication with our constructed design matrix is the best-case scenario of performance of the interpolation method.

Classification of voxels is a more direct attempt to obtain a layer volume distribution, with the property that all entries are binary, with exactly a single 1 in each row. Hence, by definition, the columns are orthogonal, and the average of the multiplication of an orthogonal design and the data is identical to regression of the data onto the same design. Therefore, classification can be viewed as a form of regression, but with a simplified design matrix.

In the limit of infinite resolution, each voxel would fall into exactly one layer and it can readily be seen that all methods would be rendered equivalent. A similar scenario presents itself when the cortex is exactly aligned with the layering and each voxel falls into precisely one layer. The aforementioned methods have been implemented in a variety of ways. Hence, the benefit of their descriptions in a consistent framework allows for easy comparison throughout the rest of this paper.

## 3 Methods

The performance of our layer extraction method is assessed by means of three experiments. First, we test the principles of the method on a simulated cortex. The model cortex has physiologically acceptable folding parameters and its layering satisfies equivolume conditions. An OLS estimation is used, as there is no (un)correlated noise added to the system. Secondly, to get a detailed understanding of the behaviour of the spatial GLM with a high number of layers, we used high resolution (post mortem) data from the primary visual cortex (V1). V1 shows a particularly strong layer structure due to the highly myelinated layer IVc (stripe of Gennari), such that the comparative performance of the methods could be easily evaluated. Thirdly, as the method is likely to be used on human in vivo data, we subsequently assessed anatomical profiles for 11 subjects. We give a detailed account of the influence of the extracted number of layers and we investigate the performance of different FWHMs that can be used for a GLS estimation. All layerings were performed on upsampled data of twice the resolution. As previously described, the best-case scenarios of the other two methods can be easily characterised in the same theoretical framework as our proposed method. Hence, in order to make the cleanest comparison between methods, all extraction methods start from the same layer volume distribution.

- The GLM method: The layer volume distribution is used as design matrix and regressed against the voxel signals.
- The interpolation method: the same design is used, but normalised (division of each element by the sum of its column) and multiplied with the data instead of regressed.
- The classification method: a regression is used, but the layer presence in the design is redistributed per voxel in a winner-takes-all manner.

### 3.1 Model cortex

In order to most cleanly compare the different methods, we established a gold standard for cortex layering. We simulated a cortex as a spring-mass system, capturing the key properties of the cortex. Most importantly, as mentioned above, the cytoarchitectonic layers of the cortex approximately conserve volume ratio over sulci and gyri, which has become known as Bok’s principle [26, 27] and is implemented in CBS Tools [43]. The intention of the simulation was to generate a layered model cortex that is consistent with the underlying assumptions of the layer extraction methods, rather than to generate a fully physiologically plausible model of the cortex. The equivolume principle leads to the best description of cytoarchitectonic layering available to date, but still does not precisely capture the layer locations [44].

Six initially equi-distant layers were generated in a two-dimensional piece of cortex that was positively and negatively curved, to simulate gyri and sulci respectively. Note that these layers are not intended to be equivalent to cytoarchitectonic layers. The layers started out with unequal volumes but were allowed to evolve until the volumes of all layers were equal, up to a precision of three orders of magnitude smaller than their size. This is illustrated in Figure 2. A detailed description of the simulation is outlined in Appendix A.

**Fig 2.**
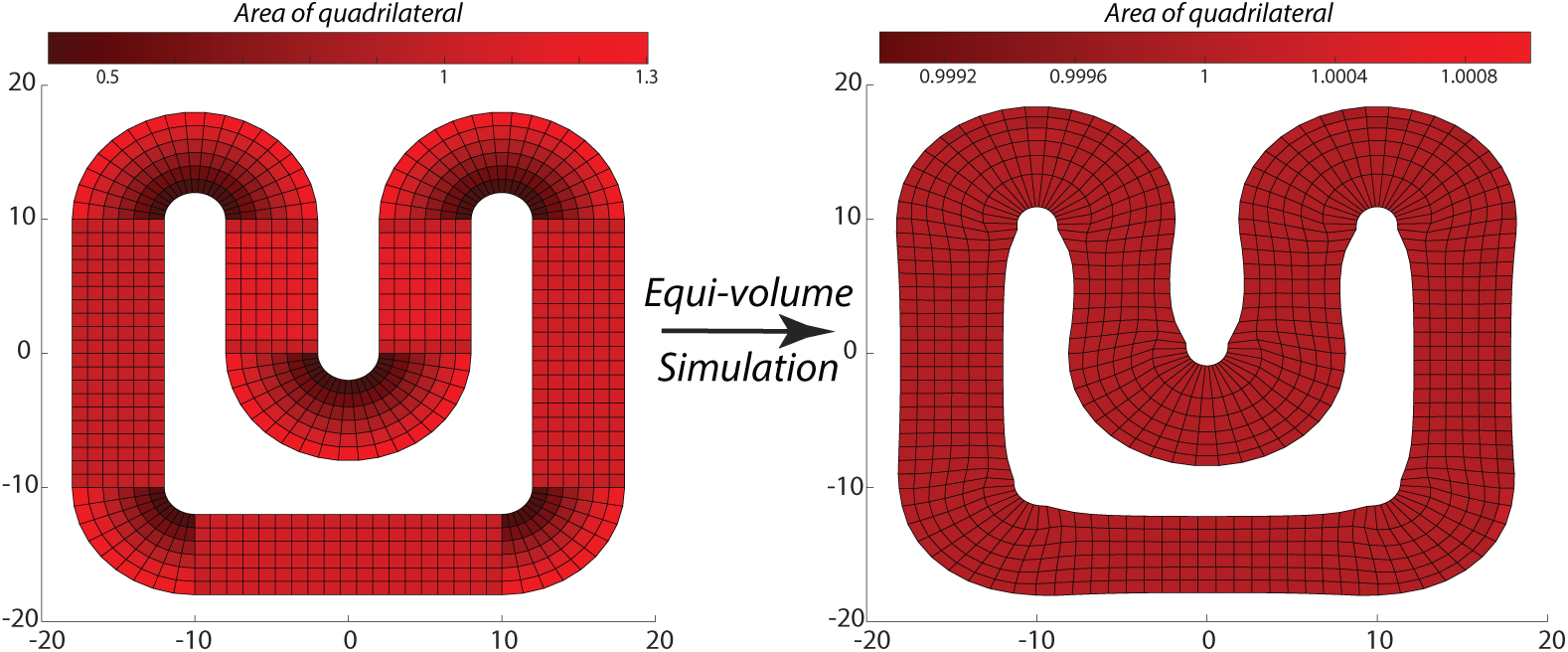
From an initially equidistant mesh, on the left, we let the points on the mesh rearrange itself in an equivolume manner. The area of a single quadrilateral is indicated by its colour. The resulting mesh, on the right, is rearranged such that all quadrilaterals had unit area (±10^-3^). Note that as a result, the layers start varying in thickness in the inner and outer bends.

The two-dimensional simulation was first rotated to break alignment with the voxel grid. Next, it was extruded to the third dimension, and resampled to a 64^3^ voxel grid. The simulation covered approximately one voxel per layer. With six layers and an approximate cortical thickness of 3.0 mm [32, 33], the volume mimicks a resolution of [0.5 mm]^3^. The outer boundaries from the simulation, corresponding to the white matter and pial surfaces, were taken as input for the layering methods. The cortex was divided into six layers and the layering was performed on upsampled data, a factor 2 in each dimension. Treating the simulated layers as a gold standard, the signal leakage between layers can be determined in terms of a spatial point spread function (PSF). The PSF of all methodologies is determined by simulating volumes in which one layer is given the value one; the remainder are set to zero. The extent to which this single layer signal can be retrieved in the correct layer is represented as a PSF. This analysis was performed on a small part of the simulated cortex (ROI shown in Figure 3) such that positively and negatively curved regions were equally represented. In order to investigate the effect of spatial resolution on the PSF, the simulated data of [0.5mm]^3^ resolution was downsampled to [1.0 mm]^3^. The same boundaries and the same layering methods and signal extraction procedures were used. No noise was added to the data, so likewise, we did not model any noise covariance in the regression equation and used the ordinary least squares solution.

**Fig 3.**
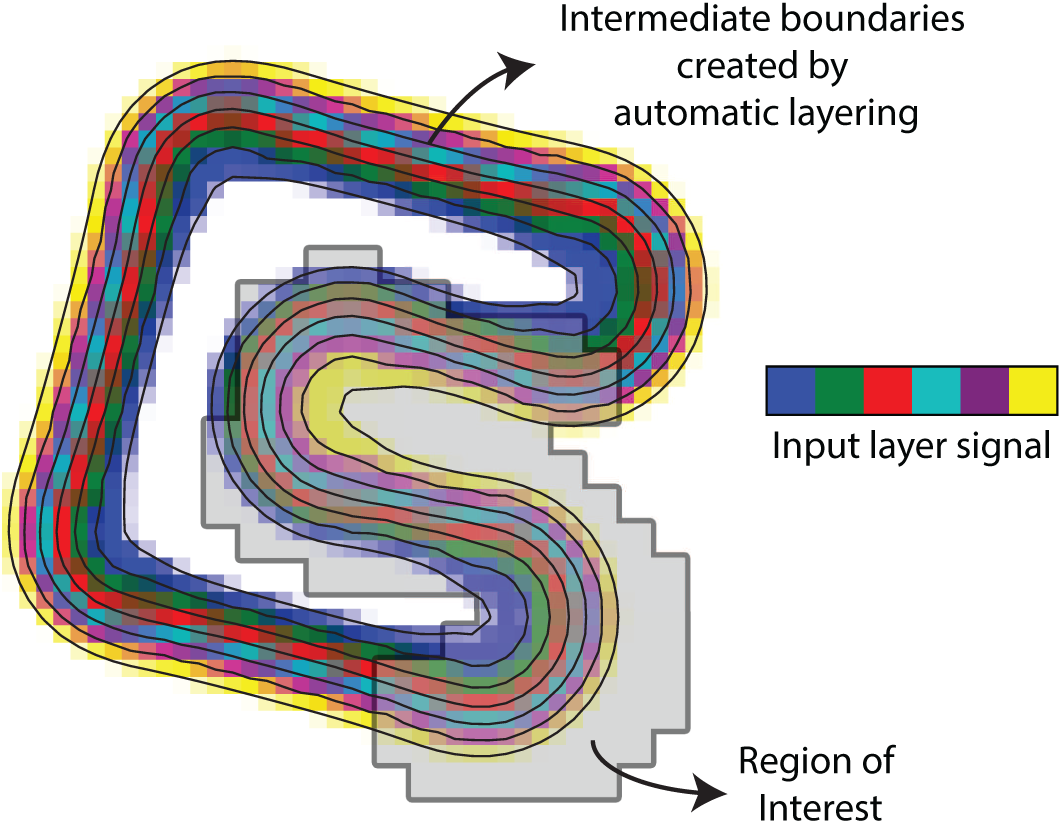

### 3.2 High-resolution data

In order to assess the performance of the layer extraction method, we examined a high-resolution post-mortem sample of the visual cortex (V1) of [0.1 mm]^3^ isotropic resolution. The thickness of this particular region was measured to be between 2 mm and 3 mm. The full experimental setup has been described by Kleinnijenhuis et al. [45]. Briefly, prior to MR imaging, samples were fixed (>2 months), soaked in phosphate buffered saline (>72h) and mounted in a syringe with proton-free liquid (∼24h). An MGE (multiple gradient-echo) sequence was used with parameters: TR=3.2 s; 7 echoes; TE1=3.9 ms; echo spacing=5 ms; matrix=250×180; FOV=25×18 mm; TA=612 s. The echoes were averaged and bias field corrected.

The aim was to extract anatomically accurate profiles including the stria of Gennari, a myelinated band of nerve fibres running parallel to the surface that is clearly visible in the image. We wanted to investigate the comparative performance of all methods in a real human cortex, but on clean high resolution data. This way, there was a clear image of the true profile, and sufficient detail that should be revealed in the extracted profile. We classified the grey matter by means of thresholding and manually adapted it to ensure accuracy over the entire region of interest. The pial surface and the white matter surface were created based on these segmentations. From these boundaries, the level set was computed and the layering was performed with 20 equivolume layers. Three regions of interest were taken, shown in Fig 6. They varied in curvature and respectively contained 1757, 924, and 1246 voxels and were 2.07 mm, 2.09 mm, and 1.97 mm thick, so this is equivalent to one layer per voxel. The results were qualitatively compared.

### 3.3 MP2RAGE data

Lastly, the method was applied to extract profiles from in-vivo data. We examined the cortical profile of the calcarine sulcus in 11 subjects from a T1-weighted MP2RAGE, acquired with a Siemens 7T scanner, with an isotropic resolution of 1.03 mm^3^, TR/TE/TI1/TI2 = 5000ms/1.89ms/900ms/3200ms, of the calcarine sulcus. The MP2RAGE was chosen for its homogeneous contrast and sharp transition from white to grey matter, such that the leakage effect to neighbouring layers could be investigated. All scans were processed by FreeSurfer [46] by means of recon-all and the boundaries generated were used in our layer pipeline. We investigated the effect of number of layers by segmenting the volume into 2, 4, 6, and 8 layers. Additionally, we wanted to test the assumption of correlated noise that we proposed in order to use generalised least squares. We compared four different FWHMs for the noise covariance, 0, 1, 2, and 3 mm, where the 0 mm effectively reduces to an ordinary least squares solution. The region of interest was a small portion of the V1 label from the Destrieux atlas that is automatically generated by FreeSurfer [47]. It was trimmed to a small part around the calcarine sulcus, because a fundamental assumption of the GLM is that the layer signal estimates are identical across the entire cortex. This cannot be guaranteed over large patches of cortex, especially because it is known that the myelination throughout the visual cortex is variable, e.g. higher around the calcarine sulcus [48]. The number of voxels in the ROI was 2009 ± 494 (*µ* ± *σ*) and the average thickness was 3.4 mm ± 0.3 mm (*µ* ± *σ*). An example for a representative subject is shown in Figure 4. The profiles were extracted on the same volume on which the segmentation and cortical reconstruction were performed, so there was no need for image registration.

**Fig 4.**
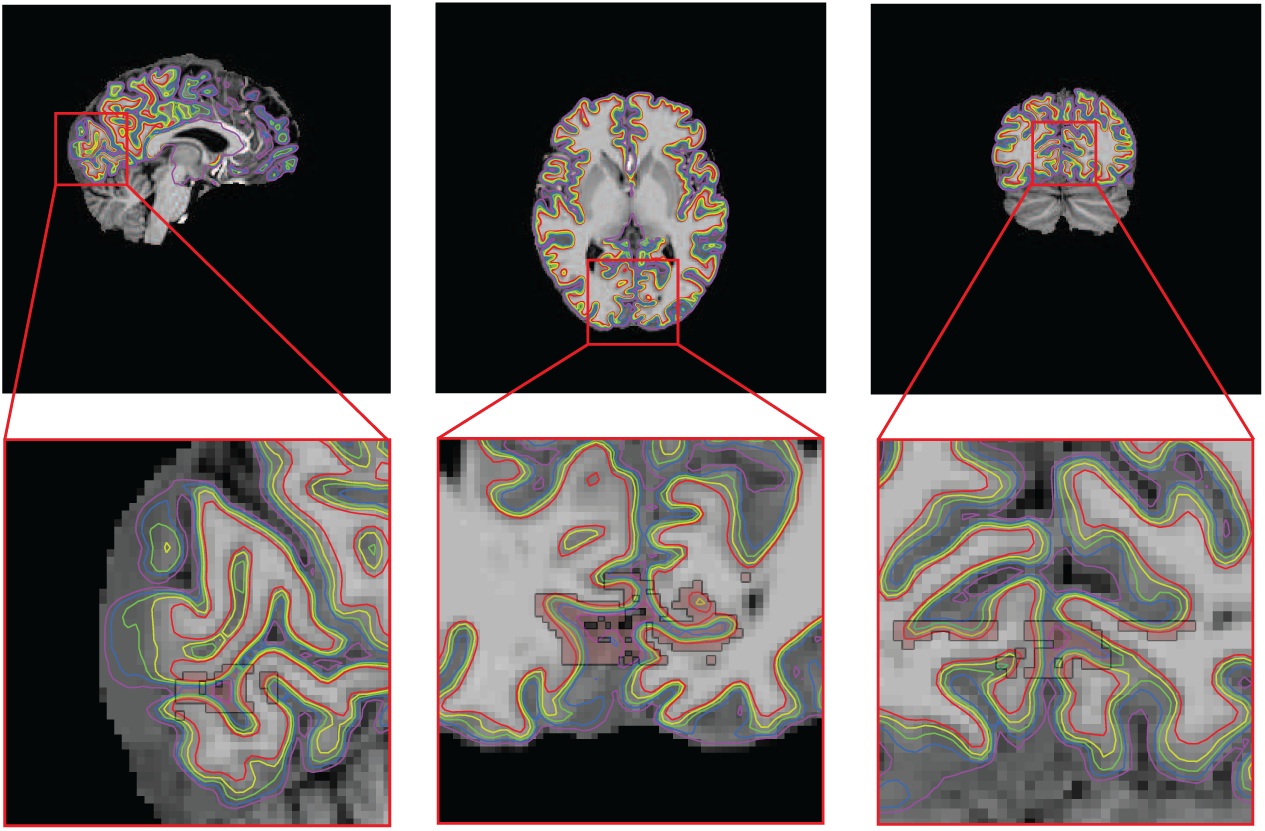
The layering (rainbow colours) and the region of interest (pink) for a representative subject. A small portion of an anatomically defined V1 region was taken in order to investigate the cortical profile in the region.

## Data Availability

All source code for the spatial GLM is freely available at https://github.com/TimVanMourik/OpenFmriAnalysis under the GPL 3.0 license. The respective modules are also available in Porcupine https://timvanmourik.github.io/Porcupine, a visual pipeline tool that automatically creates custom analysis scripts [49]. All code to generate the images in this paper are available at the Donders Repository https://data.donders.ru.nl/login/reviewer-27893111/mzQG7aE5xx2F0zOT1U0qmyVS1hC45dADMqqDWUqn3TU.

## 4 Results

We here show the results of a cortical layering on simulated data, human ex vivo data, and human in vivo data. We compare the extracted laminar signals for three different methods, the GLM, interpolation, and classification approach.

### 4.1 Model Cortex

The three different layer extraction methods were first applied to the modelled cortex, in order to estimate a point spread function of the method in ideal circumstances. The layer profiles of all layers were aligned and averaged and are shown in Figure 5 for both resolutions ([0.5 mm]^3^ and ([1.0 mm]^3^). The full unaveraged point spread functions are also shown in Supplementary Figure S1 in matrix form. The ideal PSF is a single peak of height one at the origin with no leakage to neighbouring layers.

**Fig 5.**
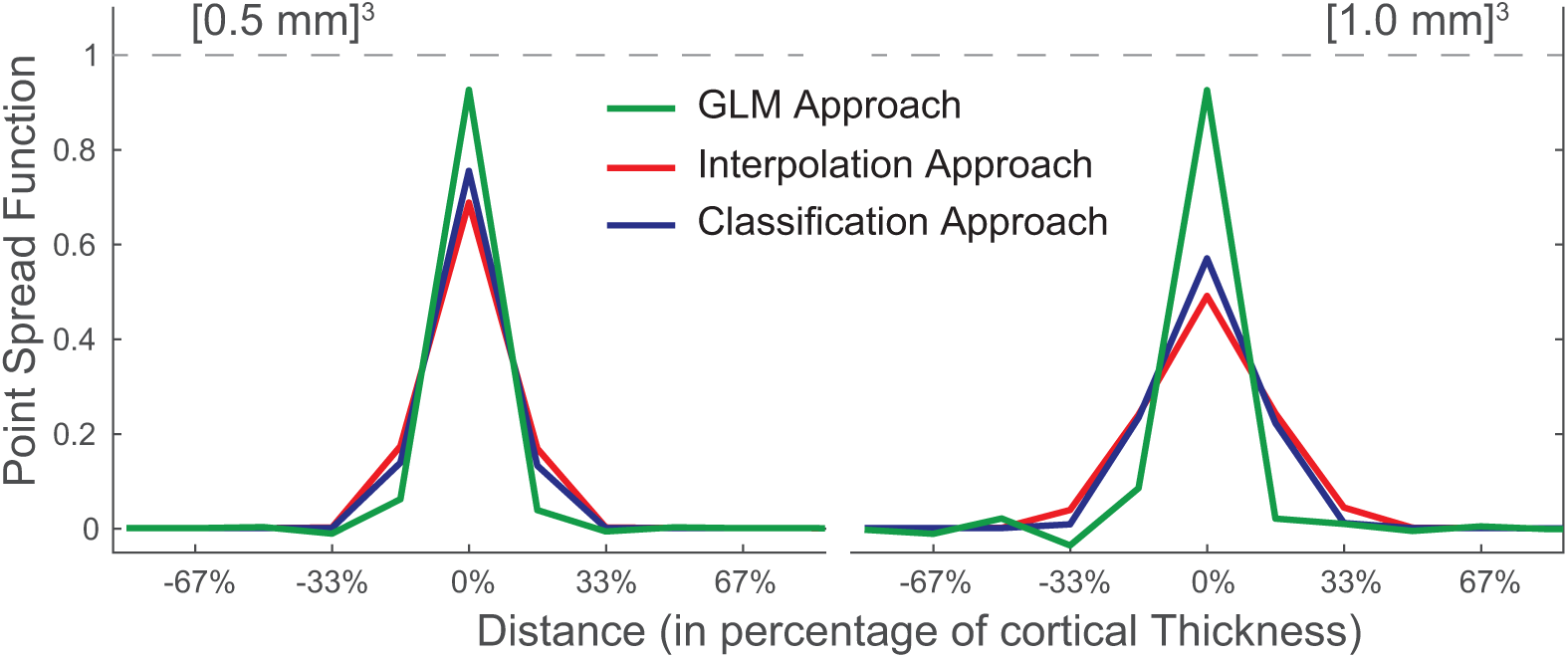
The performance of the three different approaches of obtaining layer signal, represented as a point spread function (PSF) obtained on simulated data. An ideal PSF would be an unit peak at the origin. The results are shown for approximate resolutions of 0.5 mm and 1.0 mm, on the left and right respectively. The GLM approach has a sharper PSF and is able to retrieve more signal, but potentially at the cost of a small undershoot in neighbouring layers.

For the 0.5 mm resolution volume (i.e. one layer per voxel), the peak of the distribution for the GLM reaches 92,5%, which is considerably higher than the 75.4% for the classification approach and 68,7% for the interpolation approach. This means that the latter two approaches respectively lose approximately a quarter and a third of the signal to neighbouring layers, as opposed to a only 7.5% in the GLM approach. For all methods, the leakage is close to symmetrical. The small remaining asymmetries are likely to be related to a small imbalance in proportion of voxels with a positive and negative curvature.

Also for the 1.0 mm scenario, the PSF for the GLM approach is considerably sharper. The GLM approach peaks with 92.4%, the classification approach with 56.9%, and the interpolation approach with 49.0%. As expected, the PSFs for the interpolation and classification approach are less sharp for coarser resolutions. Surprisingly, the GLM approach peaks higher, but this comes at a cost: several undershoots are visible in a sinc-like oscillating pattern. Effectively, this artificially boosts the peak signal by ‘stealing’ it from other layers. The spatial design matrix is more ill-conditioned as the number of layers is double the number of voxels over the thickness of the cortex.

The same analysis was repeated without including the gradient estimate in the layering, and instead using a cubic polynomial approximation for the partial volume kernel [23]. The resulting PSFs were identical up to 2% margin, showing that incorporating this extra type of prior knowledge has merely marginal effects on the outcome.

### 4.2 High resolution data

The extracted profile of the high resolution data is shown in Figure 6 together with an image of the data in which the region of interest is delineated. The structure of the cortex is clearly visible in the extracted profiles. It shows the intensity difference around the stria of Gennari. Additionally, towards the pial surface there is a drop in intensity of which the anatomical origin is unknown. Also note the sharp transition at the pial boundary, quickly dropping to almost zero. The average profiles look like accurate reflections of the ROI, but all methods performs roughly the same. It should be noted, however, that in all regions the GLM shows some oscillating behaviour which is likely to be artifactual to the method. This can easily be related to the sinc-like point spread function that was computed in the simulation. This effectively represents a kernel that is convolved with the true profile and thus shows the same oscillatory behaviour, much related to Gibbs ringing [50]. In particular, the artifacts proliferate at the edges of the cortex, as they scale as a function of the differences between neighbouring layers.

**Fig 6.**
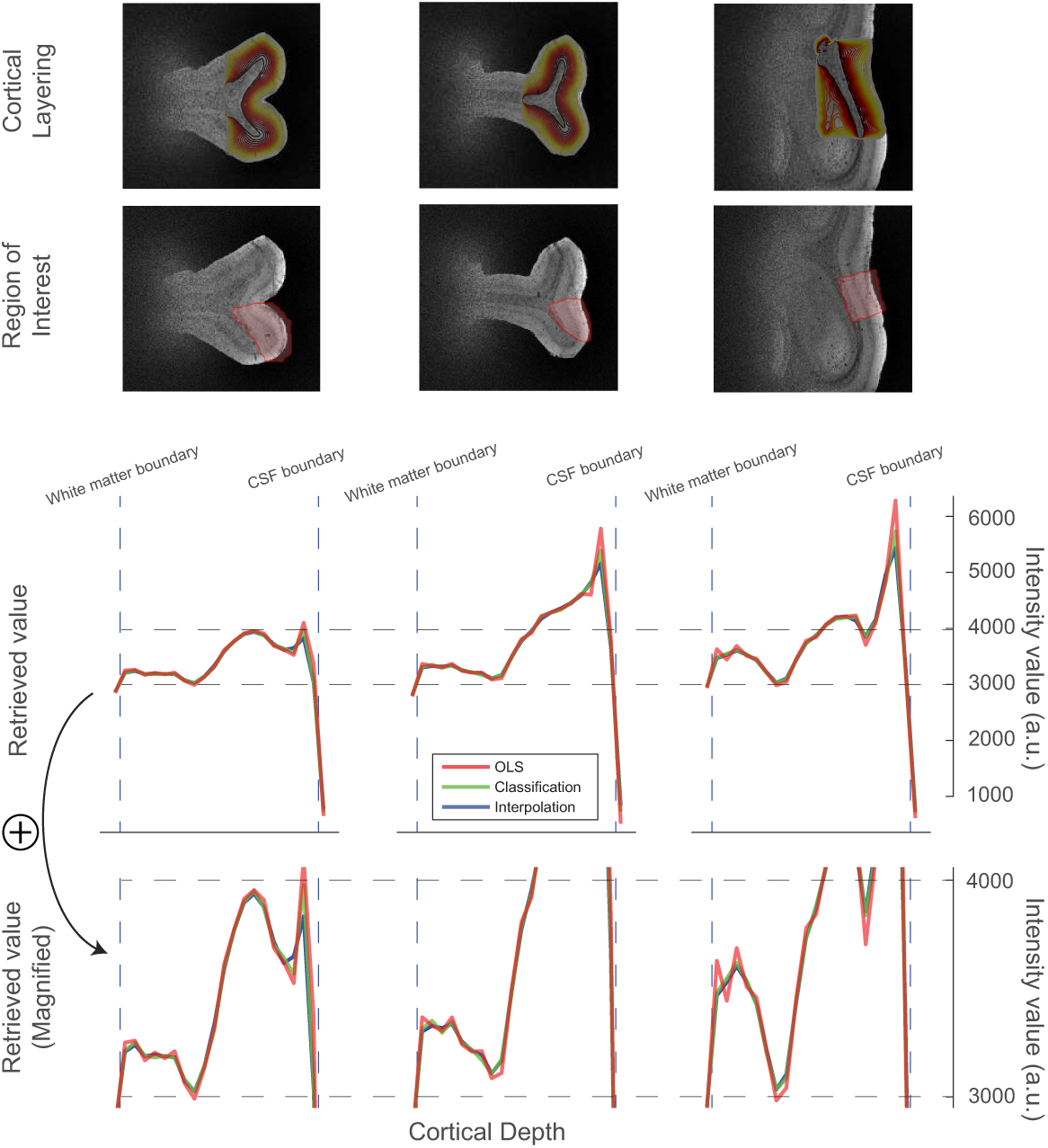
The layering, the regions of interest, and the extracted cortical profiles for three regions. While all three methods perform almost identically, there are small oscillations present in the profiles as produced by the GLM. Especially in the top layer, the peak is potentially mistakenly higher than both other methods suggest.

### 4.3 MP2RAGE data

The cortical profiles of the primary visual cortex for 11 subjects is shown for a variety of methods in Figure 7. First, the three main methods were compared based on the average over subjects. The error bars represent the standard error of the mean. The classification and interpolation approach both show smooth monotonically decreasing profiles for any number of layers. In all case, the GLM method estimates the WM signal to be higher and the CSF signal to be lower than both other methods. This could reflect a lower partial volume leakage to neighbouring layers, but may be indistinguishable from an edge enhancing artifact similar to the ones visible in the previous results. Without a gold standard, this cannot be assessed. In contrast to the two other methods, the GLM starts showing oscillating behaviour when the cortex is divided into more layers. In particular, the artifacts seem to increase dramatically when the number of layers is higher than the number of voxels. While the average over subjects is still relatively smooth, the increasing standard errors already suggests higher subject specific differences. This is especially visible in the subject specific profiles (second row of Figure 7). The highly fluctuating individual profiles for 8 cortical layers is unlikely to reflect any true underlying anatomical variation. In general, no method seems to be able to extract anatomical details, such as the stripe of Gennari. Anecdotally, the stripe is visible in some subjects, but it does not survive the anatomical variation in combination with the sensitivity limitations of the layer extraction pipeline.

**Fig 7.**
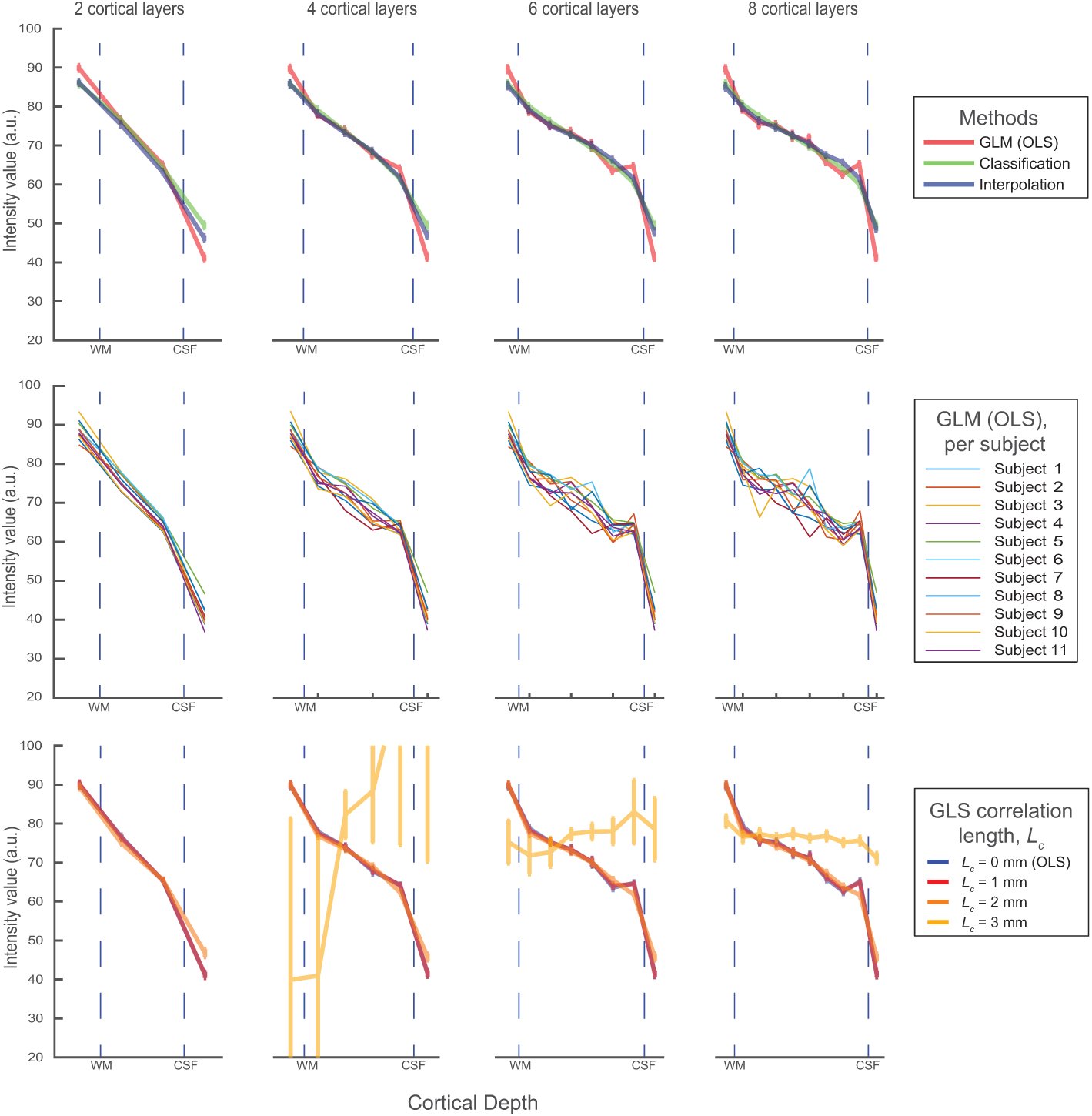
The obtained profiles for a small piece of the primary visual cortex, based on 11 subjects, for a varying number of layers (columns). In the first row, the three different methods are compared. The second row shows the individual profiles for the GLM method, showing that the solution becomes unstable when higher numbers of layers are used. In the bottom row, different correlation lenghts are tested in a generalised least squares solution.

Lastly, we investigated the assumption of correlated noise in the volume. We varied the correlation length of an assumed Gaussian noise correlation, performed a generalised least squares regression, and investigated the average profiles. For *L*_*c*_ = 0 mm, the solution reduces to an ordinary least squares problem. It can be observed that for a small correlation length (1 mm), there is only a marginal difference with no correlated noise at all. With a larger correlation length (2 mm), all profiles become somewhat smoother, but for larger values (3 mm) results start to wildly fluctuate to the extent that they are uninterpretable. It can therefore be concluded that GLS should only be used with extreme care, and that the results with the tested covariance matrices show marginal improvements at best over OLS.

## 5 Discussion and Conclusions

In this study, we propose a new method to reduce the inherent blurring of laminar profiles of current methods. Instead of interpolating a volume and averaging over a region of interest, we propose to unmix the laminar signals by using a spatial General Linear Model (GLM). In order to further reduce partial volume contamination we propose using the orientation of the voxel with respect to the cortex to better model the layer contributions to each voxel. While this provides an additional type of prior knowledge to incorporate into the layer estimation, the improvements on the layer estimates are marginal. We compute a spatial Point Spread Function (PSF) of existing cortical signal extraction methods on simulated data and explore the benefits and caveats of the spatial GLM when it is performed on human structural MRI data. On simulated data, we show that the GLM clearly outperforms existing methods, especially on a coarser resolution. However, it may be more sensitive to the imperfections of real human MRI data and result in artifacts in the extracted profile, mainly when a high number of layers is used. An initial version of this method has been applied to functional data by Kok et al. [12] and in Van Mourik et al (2018, in prep [36]).

The framework of the GLM is a well described mathematical tool and many principles transfer directly to our proposed spatial application. The core assumption of the GLM (as well as of existing methods) is that the laminar signals across every layer within the ROI are assumed to be constant. This means that any bias field that stretches through the region of interest may be detrimental to the results. Another important assumption is the normality of errors, either uncorrelated in an ordinary least squares estimation, but potentially correlated for a generalised least squares estimation. This normality is not guaranteed (and sometimes not even expected) in a laminar GLM, due to the many different sources of noise. Apart from thermal noise in the data, important sources can be the presence of e.g. blood vessels that systematically bias some part of the region of interest. At least as important as noise in the data, is noise in the model. Whenever the layer specific design matrix does not match the true underlying structure, (systematic) errors are likely to appear. While the assumed equivolume model for the cortex is the best description to date, it cannot be assumed to be a flawless description of the true cortical layering. Additionally, algorithmic implementations by necessity make numerical approximations that may induce noise as well. Correct layering also depends on the quality of the cortical reconstructions that may contain errors, especially in regions where the cortex is thin (i.e. primary visual or somatosensory cortex), highly myelinated (i.e. primary areas), or regions of reduced signal (e.g. temporal lobe, but highly dependent on acquisition). Related to this, there is a high co-occurence of neighbouring layers in the same voxels, which directly translates into a high covariance between neighbouring layer regressors. In general, covariance between regressors may induce anticorrelations, closely related to the well known anticorrelations found after global signal regression [51]. It should therefore come as no surprise that we find the point spread function of the GLM to have sinc-like characteristics and that profiles with many layers (i.e. more heavily correlated regressors) show oscillating patterns.

It is well known that a temporal design matrix needs to be balanced over conditions. Conditions need to be represented equally in the model, or otherwise the estimation may be biased towards overrepresented conditions. Similarly, it is important to have a balanced spatial design. If not, the estimation will be biased towards the overrepresented layer. This has an immediate practical implication: our implementation allows for differing layer thicknesses, which can be useful in order to match the cytoarchitectonic layer thickness. But care must be taken, as this may introduce a bias towards the thicker layers as they contribute more to the squared error. We do not provide error margins on our retrieved layer estimates, as the number of degrees of freedom in our data is not equal to the number of voxels. A valuable course for further research could be a more accurate estimation of the true degrees of freedom in order to get a better handle on the reliability of the extracted layer profiles.

The main caveat of the GLM method is the potential anticorrelation that is artificially induced in neighbouring layers. This artifact presents itself in space, but also directly translates into lower temporal correlations between neighbouring layers. As a result, one may easily conclude that neighbouring layers are temporally more distinct than is justified. Additionally, this artifact is amplified when the difference between neighbouring layers is large. Unfortunately for fMRI, this is mainly at the white matter boundary and the CSF boundary, and consequently primarily affects the deepest and highest layers. A hypothetical equal activation over the cortex may thus be amplified to appear like deep and top layer activation. If an odd number of layers is chosen, effects from both sides may even amplify to push down every second layer. While an unmixing model alludes to a superresolution potential, we strongly advise against using it as such. Using more than one layer per voxel may compromise the stability of the extracted signals. This is also illustrated by initial use in Huber et al. [13] where significant noise enhancement is observed compared to other methods.

An interesting extension of our proposed spatial GLM could be a more seemless integration with a temporal GLM, analogous to the commonly performed first and second level analysis. This spatio-temporal regression is currently performed as a two-stage approach, but could also be combined in the form of a mixture model. This is more powerful due to reduced propagation of errors [52] and would directly yield task-specific laminar results. As we here focus on the validation of the single time point scenario, this is outside the scope of this paper. A different line of improvement could be a more bottom-up approach with a forward modelling perspective of the same problem: a perspective where hypothesised laminar signals is multiplied with the layer model and compared to measured data. We here took the top-down approach by taking an existing mathematical framework, but experienced artifacts in the result as a consequence of the model inversion. Building this up in a different mathematical context may get around these violations of assumptions and provide a formulation that is closer to the problem at hand. Integrating a spatial component into a temporal layer specific hemodynamic forward model [53] could be a interesting starting point.

Hitherto, a mathematical framework has been lacking which has made it difficult to assess certainty estimates of laminar signals, which in turn has made it difficult to apply rigorous statistics. With this work, we hope to provide a contribution to such a framework in the field of laminar (f)MRI, such that it can be conducted on a more routine basis. The main use of this technique is envisioned in fMRI, where better layer extraction will allow a closer examination of layer specific BOLD in functional MRI. This may give new insights regarding feedback and feedforward connectivity of cortical areas. The spatial GLM poses improvements to dealing with the partial volume effect and prevents leakage to neighbouring layers. While there are several caveats of applying the spatial GLM on real data, we show that the performance on simulated data is far better than existing methods. We thus suggest that the price paid for a higher accuracy in ideal data is a higher susceptibility to less than ideal data.

## 6 Acknowledgements

We thank Michiel Kleinnijenhuis for providing the high-resolution data. We would like to thank Daniel Gallichan, Martin Havlícček and Ron van den Burg for the helpful discussions. Tim van Mourik acknowledges support by the Spinoza grant [SPI 40-118].

## A Spring-Mass System

### A.1 Introduction

The cerebral cortex consists of a convoluted surface of gyri and sulci. There are accurate models for the gyrification of the cortex as a whole [54] and there is a description of how the layers within the cortex behave [26]: the volume ratio between layers is equal for an arbitrarily curved pieced of cerebral cortex. This is what has become known as the Bok-principle. Here we describe how to simulate data for a two dimensional system that obeys Bok’s principle by means of a spring-mass system. A spring-mass system was chosen, because it is independent of the algorithms by means of which we estimate curvature in the volume. This makes it ideal benchmark data for the methods presented in the body of this paper.

## A.2 Spring-Mass System

The cortex is modelled by means of a spring mass system. This is an approximation of the cortex consisting of quadrilaterals. Each quadrilateral consists of four edges and four vertices, which are the springs and masses respectively. The collection of springs and masses that form quadrilaterals will henceforth be referred to as the system.

The system has total energy *U*. In the present simulation only a single contribution to the energy is considered, related to the area of each quadrilateral. More generally, other contributions could be taken into account, for example relating to the length of each edge:

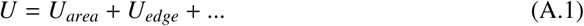

We are looking for the case where the energy is minimised, as this is when the system has come to rest and all quadrilaterals have reached an equilibrium area. Note that only the vertices are displaced in the first instance; the edges and quadrilaterals are formed as a consequence. The energy decreases by moving the vertices in the direction of the net force applied to them. The force 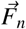 on vertex *n* is minus the derivative of *U* with respect to the position 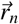 of that vertex:

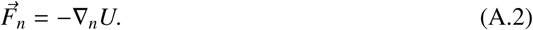

A quadrilateral *Q*_*i*_ is defined to be the space enclosed by four vertices in two dimensions 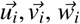 and 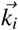, shown in Figure H. Its area *A*_*i*_ is a scalar function depending on the vertices of the quadrilateral, 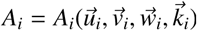

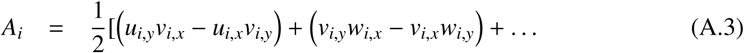

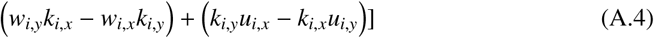

**Fig H.**
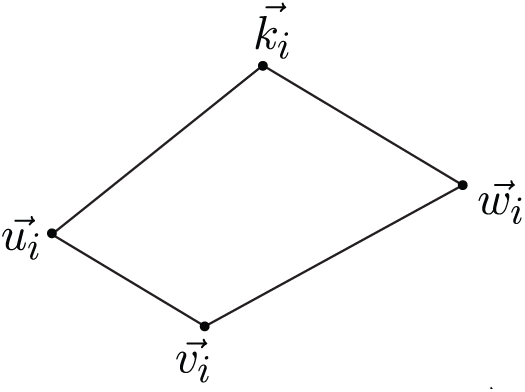
Quadrilateral *Q*_*i*_, consisting of vertices 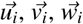 and 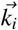.

The vertices 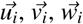 and 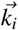 are chosen such that *A*_*i*_ *>* 0.

Let *Q* = {*Q*_1_, *Q*_2_, *Q*_3_, …} be the set of all quadrilaterals. The energy of the system is

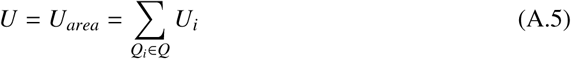

where *U*_*i*_ is the energy of quadrilateral *Q*_*i*_

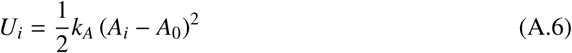

such that the quadrilateral energy is a function of the difference between the actual area *A*_*i*_ and the equilibrium area *A*_0_.

Subsequently, the gradient is computed for the energy stored in the system:

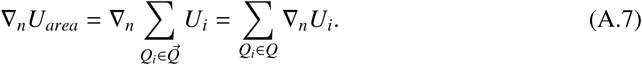

The gradient of a single quadrilateral energy term takes the form:

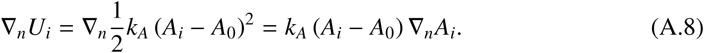

Note that ∇_*n*_*U*_*i*_ can be non-zero only if vertex *n* belongs to *Q*_*n*_, i.e. if 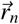 is one of the vertices 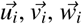 or 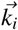. To be explicit, let 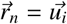. In two dimensions 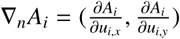. The two components follow immediately from A.4:

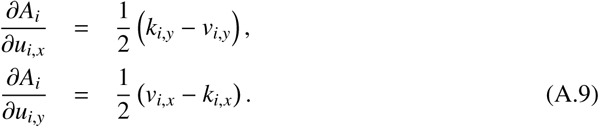

Combining equations A.2, A.8 and A.9, the resulting force on vertex *n* due to the preservation of volume is

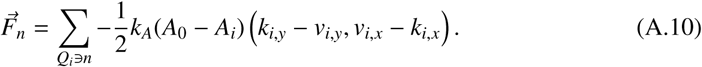

Here the summation is over the quadrilaterals *Q*_*i*_ that contain vertex *n*, and 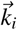 and 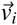 are the neighbours of 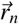 in *Q*_*i*_. The vertex will come to rest if *A*_*i*_ = *A*_0_, and the strength of the force can be adjusted by parameter *k*_*A*_. All forces are additive.

This was implemented in C++ as a stand-alone application. The program reads in a mesh and evolves the vertices until the system comes to rest.

### B Orientation dependent partial volume distribution

In order to find out how different laminae are distributed over voxels, it is important to know the orientation and location of the surface with respect to the voxels. Here we analytically describe an algorithm to solve this problem.

The intersection of a laminar surface and a voxel is approximated by the intersection of a cuboid and a plane that are arbitrarily positioned and oriented with respect to each other.

The voxel grid is given by three primitive lattice vectors 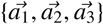, having the orientation and length of the voxel edges. The lattice vectors are usually but not necessarily oriented along the cardinal axes. For a cubic voxel grid with edge length *L* the primitive lattice vectors are {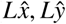 and 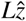}. A voxel can be indexed with three integers *m*_*i*_. The centre position 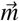 of the voxel is

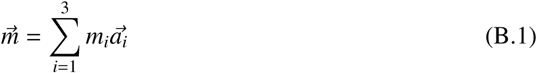

and the 8 corner positions 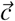 of the voxel are

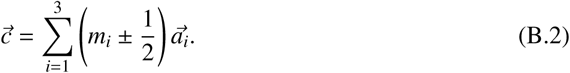

The voxel volume is 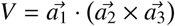

A plane can be defined by a vector 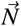 The plane is perpendicular to 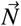 and has distance 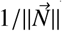 to the origin. The plane is given by all points 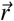 satisfying

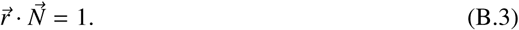

It is useful to express 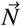 in terms of reciprocal lattice vectors 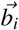:

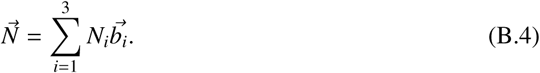

The reciprocal basis vector 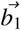 is defined as 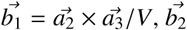 and 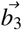 are obtained by cyclic permutation. For a cubic voxel grid, the reciprocal basis vectors are 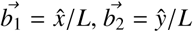 and 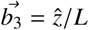. By construction 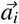 and 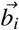 are orthonormal to each other

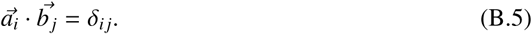

Points on the four edges of voxel 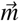 in the direction 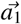 are of the form

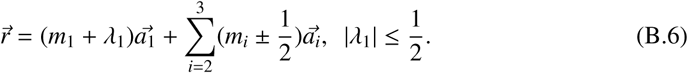

In view of B.4 the four possible intersection points of 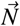 with the edges parallel to 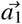 are given by the four *λ*_1_ of the form

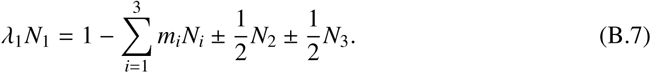

Analogously the possible intersection points with the edges parallel to 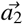 and 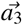 are given by four *λ*_2_ and *λ*_3_ values respectively. Of the 12 possible intersection points, those with 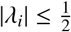 give the actual (at most 6) intersection points. The area of the polygon connecting the intersection points can be readily computed, as well as the volume on either side of the plane.

At this point we have expressions for the intersection point that are independent of the choice of the voxel edge vectors 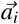. Explicit expressions can be obtained for the intersection points and intersection area. To be explicit, consider the intersection of a plane, moving with a given orientation, i.e. 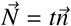 where *t* takes arbitrary real values and 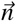 stays constant. A finite intersection is only found for the range of *t* values between the maximum and minimum value of 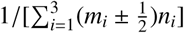

## 8 Supplementary Figures

**Fig S1.**
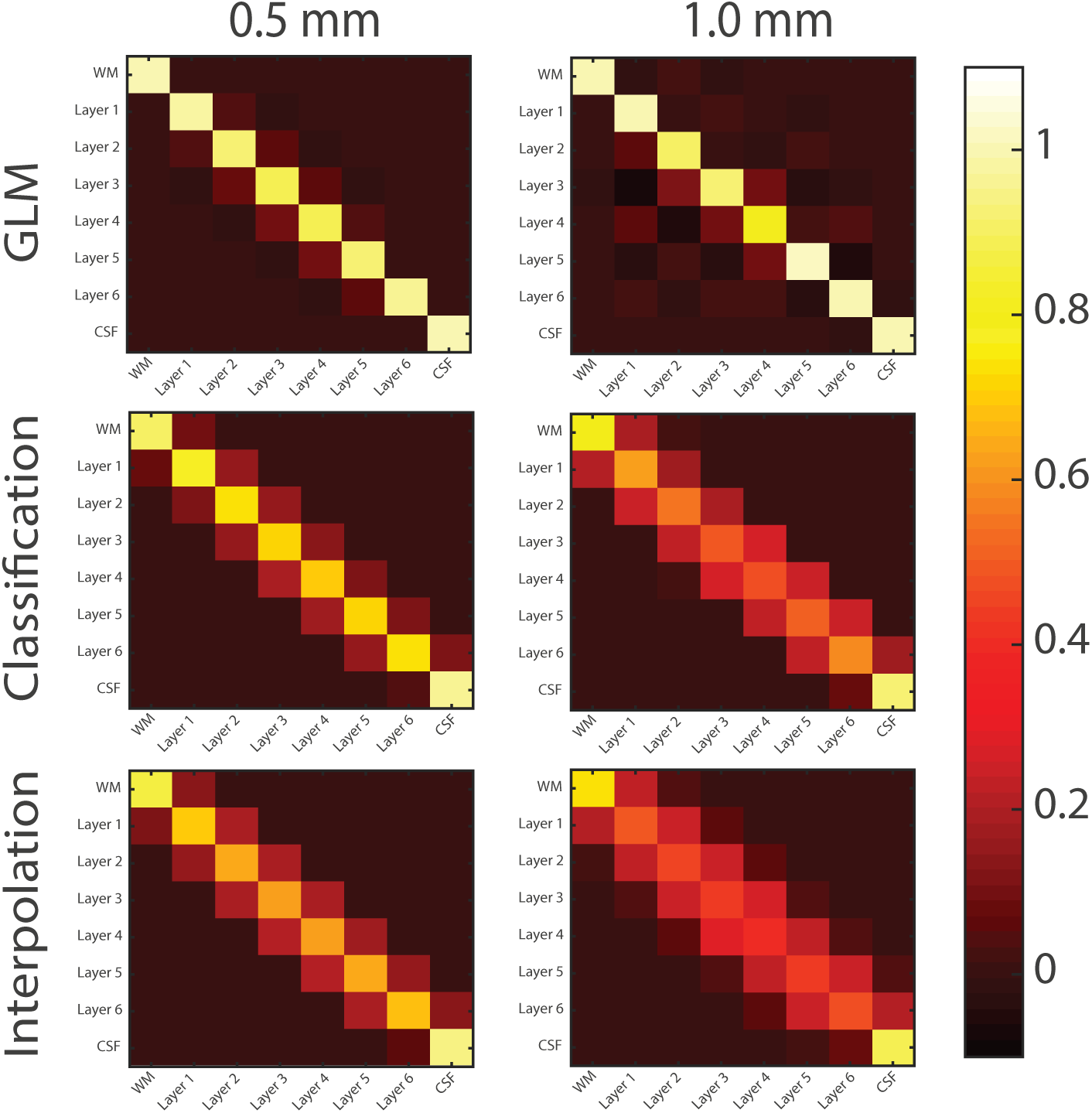
The point spread functions of all layers for both resolutions and both methods. An ideal point spread function would look like an identity matrix. This is approached more by the GLM method for both resolutions.

